# Reconciling ASPP-p53 Binding Mode Discrepancies through an Ensemble Binding Framework that Bridges Crystallography and NMR Data

**DOI:** 10.1101/2023.09.17.558183

**Authors:** Te Liu, Sichao Huang, Qian Zhang, Yu Xia, Manjie Zhang, Bin Sun

## Abstract

ASPP2 and iASPP bind to p53 through their conserved ANK-SH3 domains to respectively promote and inhibit p53-dependent cell apoptosis. While crystallography has indicated that these two proteins employ distinct surfaces of their ANK-SH3 domains to bind to p53, solution NMR data has suggested similar surfaces. In this study, we employed multi-scale molecular dynamics (MD) combined with free energy calculations to reconcile the discrepancy in the binding modes. We demonstrated that the binding mode based solely on a single crystal structure does not enable iASPP’s RT loop to engage with p53’s C-terminal linker—a verified interaction. Instead, an ensemble of simulated iASPP-p53 complexes facilitates this interaction. We showed that the ensemble-average inter-protein contacting residues and NMR-detected interfacial residues align well with ASPP proteins, and the ensemble-average binding free energies better match experimental Kd values compared to single crystallgarphy-determined binding mode. For iASPP, the sampled ensemble complexes can be grouped into two classes, resembling the binding modes determined by crystallography and solution NMR. We thus propose that crystal packing shifts the equilibrium of binding modes towards the crystallography-determined one. Lastly, we show that the ensemble binding complexes are sensitive to p53’s intrinsically disordered regions (IDRs), attesting to experimental observations that these IDRs contribute to biological functions. Our results provide a dynamic and ensemble perspective for scrutinizing these important cancer-related protein-protein interactions (PPIs).

## 1 Introduction

The tumor suppressor protein p53 is a transcription factor that activates genes involved in apoptosis and cell-cycle arrest upon the detection of oncogenic stress.^1^ p53 is a signaling hub that can be regulated by numerous proteins (involved in >1000 protein-protein interactions (PPIs)^2^). Consequence of p53 activation is majorly cell apoptosis, but can also cause cell-cycle arrest, depending on specific p53 regulators.^3^ ASPP2 and iASPP proteins are two p53 regulators that promote and inhibit p53-dependent apoptosis, respectively.^4^ Given the critical role p53 plays in tumor, mechanical insights into the reversal regulatory effects of ASSP2 and iASPP on p53 are much wanted yet are obscured by lacking of definite structural characterizations of the PPI.

So far, structural characterizations have primarily focused on interactions within the folded portions, specifically between p53’s DNA-binding domain (DBD) and the ankyrin repeats and an SH3 domain (ANK-SH3) of iASPP/ASPP2^5,6^ (Fig. 1A). The ANK-SH3 domains of ASPP2 and iASPP are highly conserved, sharing approximately 70% sequence identity and nearly identical crystal structures (Cα RMSD*∼*1.24 Å).^6,7^ Despite this high degree of conservation, crystallography has revealed that when binding to p53’s DBD, ASPP2 and iASPP employ entirely different surfaces on their ANK-SH3 domains: ASPP2 primarily utilizes the SH3 domain to bind p53, while iASPP employs a flat surface formed by the parallel ank-repeat helices to interact with p53 (Fig. 1B). This significant difference in binding modes has been reported to contribute to ASPP2 and iASPP’s opposing regulatory effects on p53. ASPP2 directly displaces DNA from p53’s DBD through a competitive binding mechanism,^8,9^ whereas iASPP binds to p53, triggering a molecular switch in the p53 L1 loop, interfering with p53’s DNA binding.^6^ While the general mechanism by which ASPP proteins alter p53’s binding capacity to various genes, thus influencing cellular lifeor-death processes,^6,10,8^ has been established, this mechanism is challenged by conflicting reports on the binding modes of iASPP-p53 interactions. Ahn et al.’s solution NMR data suggests that iASPP uses a similar binding surface on its ANK-SH3 domain as ASPP2 to interact with p53’s DBD^4^ (Fig. 1B), in stark contrast to the crystallography-determined iASPP-p53 complex (PDB 6RZ3).

**Figure 1:**
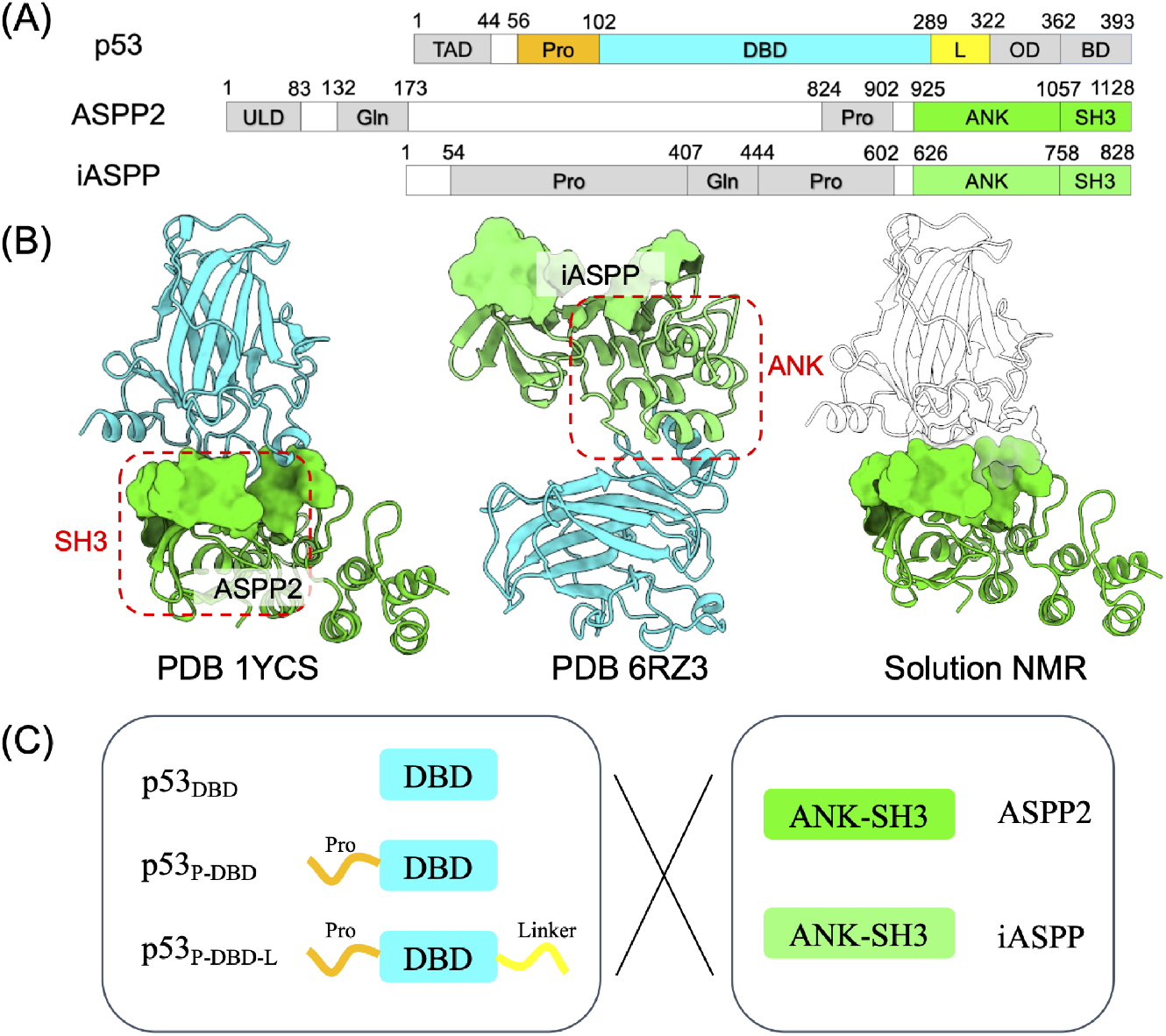
(A) Domain organizations of p53 and ASPP proteins. p53’s DBD and ASPP’s ANKSH3 are folded domains whereas the remaining sequences are intrinsically disordered. B) Crystal structures of ASPP2-p53 complex (PDB 1YCS^5^) and iASPP-p53 complex (PDB 6RZ3^6^). Solution NMR studies^4^ also suggest a iASPP-p53 binding mode that is different from PDB 6RZ3. (C) In this study, we combined all-atom and coarse-grained (CG) MDs plus free energy calculations to explore the binding mechanisms between three p53 constructs (p53_DBD_ p53_P-DBD_ and p53_P-DBD-L_) and the ANK-SH3 domains of ASPP2/iASPP.

Discrepancies between crystallography and NMR structures are frequently encountered, especially when determining biologically relevant PPIs.^11,12,13,14^ Both methods have their own advantages and shortcomings. While the structures themselves are accurate, the protein-protein complexes determined by crystallography may not be biologically relevant due to crystal packing effects.^11^ This is especially true for weak PPIs, as it has been reported that there is a greater than 50% probability that crystallography-determined complexes are not biologically relevant if the dissociation constant (K_*D*_) is greater than 100 µM.^11^ In contrast, solution NMR captures protein motion in an aqueous environment and is therefore more functionally related. However, it suffers from inaccuracies in chemical shift assignments and cannot directly provide atomistic structures; instead, it serves as constraints to derive structures that satisfy the observed chemical shifts.^12^ Solution NMR generally reflects ensemble-average properties, as demonstrated by Pochapsky *et al*, who used solution NMR to capture multiple p450 protein conformers that are all important to the protein’s function.^15^ Meanwhile, the binding mode determined by crystallography is more likely to be a subset of the ensemble of binding complexes, as demonstrated by Zuo et al., who used computational methods to show that multiple poses, including the X-ray one, are all consistent with experimental affinities.^16^ For our system of interest, whether the observed discrepancies between crystallography and NMR in determining the binding mode for iASPP-p53 are complementary or mutually exclusive is unknown and warrants further exploration.

Reconciling the discrepancy in binding modes can provide valuable insights into the roles of intrinsically disordered regions (IDRs) in regulating the ASPP-p53 PPI. NMR studies have suggested that p53’s IDRs play a role in fine-tuning the PPI. Both ASPP and p53 encompass significant portions of their protein sequences predicted as IDRs. Ahn et al. demonstrated that ASPP2 primarily binds to p53’s DNA-binding domain (DBD), while iASPP predominantly interacts with a linker region located C-terminal to the DBD domain.^4^ This highlights the influence of p53’s IDRs in distinguishing between bindings to the highly conserved ANK-SH3 domains of ASPP2 and iASPP. Furthermore, in the case of ASPP2-p53 binding, solution NMR studies conducted by Tidow et al.^8^ revealed that, in addition to the interface residues reported in PDB 1YCS, p53 residues outside the interface (such as loop 1 and the C-terminal helix) were implicated in binding ASPP2. Clarifying the relevance of these NMR-based observations to ASPP-p53 functions is expected to be possible if the aforementioned binding mode discrepancy is resolved.

In this study, we performed extensive multi-scale molecular dynamics (MD) simulations to investigate if p53’s DBD domain and ASPP2/iASPP ANK-SH3 domain could bind in multiple binding modes, and whether the controversial crystallography- and NMR-determined binding modes can be reconciled under this ensemble binding framework. We also simulated the bindings of different p53 constructs that contain extra IDRs to ASPP, to explore how p53’s IDRs finetuning the ASPP-specific PPIs. We anticipate that our structural characterizations could facilitate thearpeutic developments targeting these important PPIs.

## 2 Materials and methods

### 2.1 Conventional molecular dynamics (MD) simulations

Simulations were based on the crystal structures of p53_DBD_ in complex with ASPP2 and iASPP’s ANK-SH3 domains (PDB IDs are 1YCS^5^ and 6RZ3,^6^ respectively). The input files for MD were prepared by the tleap program from Amber20 package.^17^ The system was solvated into a 12Å margin OPC waterbox with 0.15 M KCl ionic strength, and the Amber ff19SB force field^18^ was used for protein. The system was first subject to 50000 steps energy minimization with the first 200 steps using the steepest descent algorithm and the remaining steps using the conjugate gradient algorithms. The minimized system was then heated to 300 K via a two-step procedure: 0 to 100 K in NVT ensemble over 0.1 ns followed by 100 to 300 K in NPT ensemble over 0.5 ns. During heating, harmonic constraints with a force constant of 5 kcal/mol/Å^2^ were introduced onto protein backbone atoms. After heating, a 1 ns equilibrium simulation was performed in the NPT ensemble at 300 K with reduced force constant of 1 kmol/mol/Å^2^. Three independent 1 µs long production runs (NPT ensemble, 300 K) were initiated from equilibrated system. During the simulation, all of the length of bonds involving hydrogen were restrained by the SHAKE algorithm,^19^ and the temperature was controlled by Langevin thermostat.^20^ The time step was 4 fs after hydrogen mass repartitioning^21^ and the nonbonded cutoff was 12.0 Å. Notably, the bound zinc ion in p53 crystal structures was considered in our all-atom MD simulations. This Zn^2+^ is vital to p53’s structure stability and function,^22,23^and our own test in Fig. S2 shows that removing the Zn^2+^ alters the dynamics of p53_DBD_ and changes the conformation of motifs at the PPI interface with ASPPs. However, since protein force fields generally notoriously handle polar contacts^24^, to maintain the coordination of Zn^2+^ to p53’s His83-CYS80,142,146 motif, we introduced harmonic constraints between Zn^2+^ ion and the coordinating atoms with a force constant of 300 kcal/mol/Å^2^ to retain Zn^2+^ binding. The parameters of Zn^2+^, as well as the KCl monovalent ions, were adapted from the Li-Merz dataset.^25^

Besides the p53_DBD_, we built two additional p53 constructs that have the IDR regions flanking p_53_’s DBD domain added: The p_53_P-DBD that has the N-terminal Pro-domain (residues 56-102) added, and p53_P-DBD-L_ that has the additional C terminal linker (residues 289-322) added. The initial structures of these IDRs were built from sequences using the tleap program, and were sampled from 500 ns conventional MD at 300 K in explicit solvent model. The representative structure of the most-populated cluster was selected, and was then joined to p53DBD using Pymol. Simulations of these p53 constructs’ interactions with ASPPs were started from the above mentioned PDB structures and followed the same MD protocol.

### 2.2 Martini CGMD to simulate the binding process between p53 and ASPP

The Martini3.0 coarse-grained (CG) model^26^ was used to simulate the protein-protein binding. Coarse graining of p53 and ASPP all-atom structures into Martini beads were performed using the vermouth program,^27^ and the DSSP program^28,29^ was used to detect protein secondary structure information. The bound Zn^2+^ ion in p53 was omitted for coarse graning, thanks to the elastic network strategy that can maintain protein tertiary structure. The elastic bond force constants was set as 500 kJ/mol/nm^2^ and the lowerand upperelastic bond cutoffs were 5 and 9 Å, respectively. Since long range electrostatic interactions between proteins become centrysymmetric at *∼*40 Å,^30^ the ASPP ANK-SH3 domain was randomly placed >=40 Å away from p53 construct to minimize bias. The system was solvated in a 150 *×*150 *×*150 Å cubic box with 0.15 M ionic strength using standard Martini water and NaCl models.^31^ The system was first energy minimized 2000 steps using the steepest descent algorithm and was then heated to 300 K in the NPT ensemble over 1 ns, with constraints on the protein. The equilibrated system was subject to a 4 µs production MD in NPT ensemble at 300 K using a 20 fs time step. The temperature control was achieved with the velocity rescale (V-rescale) and pressure was controlled with the Parrinello-Rahman barostat. For each system, 50 replicas of 4 µs CGMD runs were performed via Gromacs2020.4^32^ to investigate the binding between p53 constructs and ASPP ANK-SH3 domains.

### 2.3 Umbrella samplings to obtain the disassociation potential of mean force (PMF)

Umbrella samplings were performed to estimate the potential of mean force (PMF) along the disassociation pathway of the p53-ASPP complexes, starting from both crystal complex structures and Martini CGMD sampled complexes. Martini complexes were backmapped into all-atom structures using the backward.py^33^ script, and were subject to 100 ns explicit solvent MD with Zn^2+^ added to p53 to further optimize the complex following the abovementioned MD protocol. The reaction coordinate (RC) was defined as distance between centers of mass (COM) of p53 and ASPP and was binned into 0.5 Å-wide windows. Samplings in each window was enforced by harmonic constraints with 10 kcal/mol/Å^2^ force constant. Starting structures used for each window are the last trajectory frames from previous windows. For each window, a 10 ns production MD was performed in NPT ensemble at 300 K. The PMF was estimated by the WHAM program.^34^ All MD simulations performed for this work are summarized in Table S1.

### 2.4 Sketch-map projections to obtain 2D mode

The sketch-map dimensionality reduction method^35,36^was employed to project the Martini CGMD sampled ASPP-p53 complexes onto 2D plane. Each trajectory frame was encoded as a highdimensional space vector, 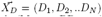 where *D*_1_, *D*_2_…*D*_*N*_ are the minimum distances between the backbone beads (Martini BB type) of p53 and ASPP. Beads were chosen from the secondary structure elements (helices and sheets) from p53 and ASPP because inter-protein distances between these rigid structures have low fluctuations compared with loops, and can thus accurately encode the binding configurations.^37^ We set N as 21: *D*_1_ to *D*_10_ represents the minimum distance of the 10 residues of p53 to 11 residues of ASPP.Namely, for residue i, we first calculated its distance to the selected ASPP residues and assign the smallest value to *D*_*i*_. And *D*_11_ to *D*_21_ are the reverse minimum distances, namely, the minimum distance between ASPP residues to p53. Projection follows the principle that points close in the high-dimensional space should be close in the 2D space,^35^ by minimizing the following stress function:^35^

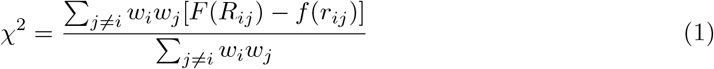

where *w*_*i*_, *w*_*j*_ are weights of points, *R*_*ij*_, *r*_*ij*_ are distances between points in the highand lowdimensional spaces, respectively. *F, f* are sigmoid functions that map distances between 0 and 1. In our calculations, points weight and sigmoid functions were taken as the default values from the sketch-map tutorial.

### 2.5 Analyses

Analyses were conducted using the CPPTRAJ program,^38^ MDTraj,^39^ and Matplotlib libraries. RMSD, RMSF and distance calculations were conducted using CPPTRAJ. 3D space distributions of p53 around ASPP were estimated using the *grid* command from CPPTRAJ. We selected a 0.5 Å spacing to build 3D grids centered on the average COM position of ANK-SH3 domain after aligning the MD trajectories on this domain, and then binned p53 densities into grids, and normalized to standard water density 1.0 g/cm^3^. Projections of MD trajectories onto 2D plane were done using the MDTraj library which aligns trajectories and extracts the Cartesian coordinate of protein atoms. Inter-protein contact data was calculated using the *nativecontacts* command from CPPTRAJ with default distance cutoff 7 Å. Structure were rendered using UCSF ChimeraX,^40,41^ and VMD.^42^ Scripts supporting this work are available at https://github.com/bsu233/bslab/tree/main/2023-p53ASPP.

## 3. Results

### 3.1 MD simulations starting from crystallography binding modes fail to capture the experiment-verified inter-protein interactions

Both NMR studies^4^ and peptide screening assays^43^ have confirmed that p53 utilizes its intrinsically disordered linker (located C-terminal to the DBD domain) to bind to iASPP’s RT loop. A designed peptide mimicking p53’s linker has been shown to competitively displace p53 from binding to iASPP, thereby blocking iASPP’s regulation of p53 in vitro.^43^ We conducted MD simulations to investigate whether p53’s linker can interact with iASPP’s RT loop in the crystal structure-resolved binding modes, aiming to justify the relevance of these modes as the biological unit. We initiated our simulations from both PDB 1YCS and 6RZ3, assuming that p53 can bind to iASPP via these two modes. The samplings presented in Fig. 2 clearly demonstrate that, starting from both crystal structure-resolved binding modes, p53’s linker is unable to reach iASPP’s RT loop. The discrepancies observed between the MD samplings and experimental findings strongly suggest that iASPP and p53 likely engage in alternative binding modes.

**Figure 2:**
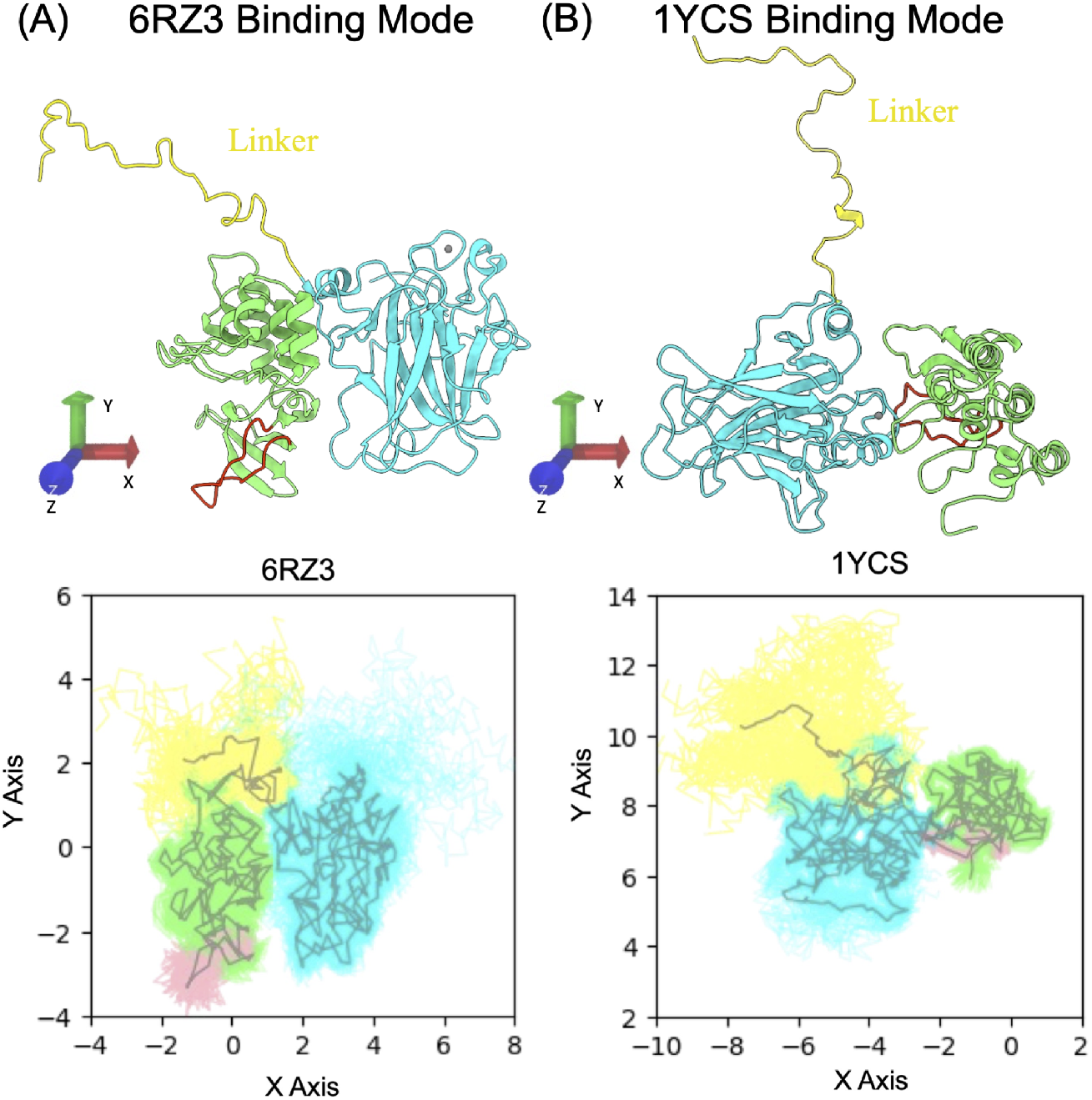
Examining the interactions between p53’s C-terminal linker (yellow) and iASPP’s RT loop (red), using 3*×* 1 µs all-atom MDs. (A) Starting from PDB 6RZ3. (B) Starting from PDB 1YCS. MD trajectories were aligned on p53’s DBD against the two reference structures, and were plotted on the XY plane. The gray lines depict the average structure of protein backbone.

### 3.2 Ensemble-average interprotein contacts agree with solution NMR revealed PPI interfaces

To explore potential alternative binding modes between p53 and ASPP, we conducted unbiased coarse-grained (CG) molecular dynamics simulations (MD) to simulate the protein-protein binding processes. We opted for the Martini 3.0 CGMD model^26^ due to its consideration of side-chain packing, which has been shown to have a critical impact on protein-protein interactions.^44^ We first simulated the binding between p53_P-DBD_ and the ANK-SH3 domains of ASPPs. Solution NMR studies conducted by Ahn *et al* ^4^ reported chemical shift changes in ASPPs residues caused by the binding of p53, allowing us to map out the interfacial residues on the ANK-SH3 domains. We therefore calculated the inter-protein contacts based on the cumulative Martini CG trajectories and compared the per-residue contacts of ANK-SH3 domain residues with the NMR data to validate our Martini CG results.

As illustrated in Fig. 3A-B, for both ASPP2 and iASPP, the per-residue contacts of the ANK-SH3 domain qualitatively agree with the NMR chemical shift data. In the case of ASPP2, the contact data suggests that the borders between ANK repeats, particularly the ANK3-ANK4 border, exhibit significant contacts with p53. Furthermore, the SH3 domain is largely involved in contacting p53. These contact patterns are consistent with the NMR chemical shift data. For iASPP, both Martini CGMD and NMR data indicate that the SH3 domain is at the protein-protein interface. Additionally, Martini CGMD reports that ANK4 is involved in interacting with p53 (Figure 3B), partially resembling the inter-protein interaction observed in the p53_DBD_-iASPP complex crystal structure (PDB 6RZ3). This suggests that the ensemble of p53-iASPP structures sampled using Martini CGMD may contain the binding mode observed in the crystal structure (which will be discussed in detail later). Overall, we observed that the binding complexes obtained from Martini CGMD qualitatively capture the protein-protein interface reflected in the solution NMR data.

**Figure 3:**
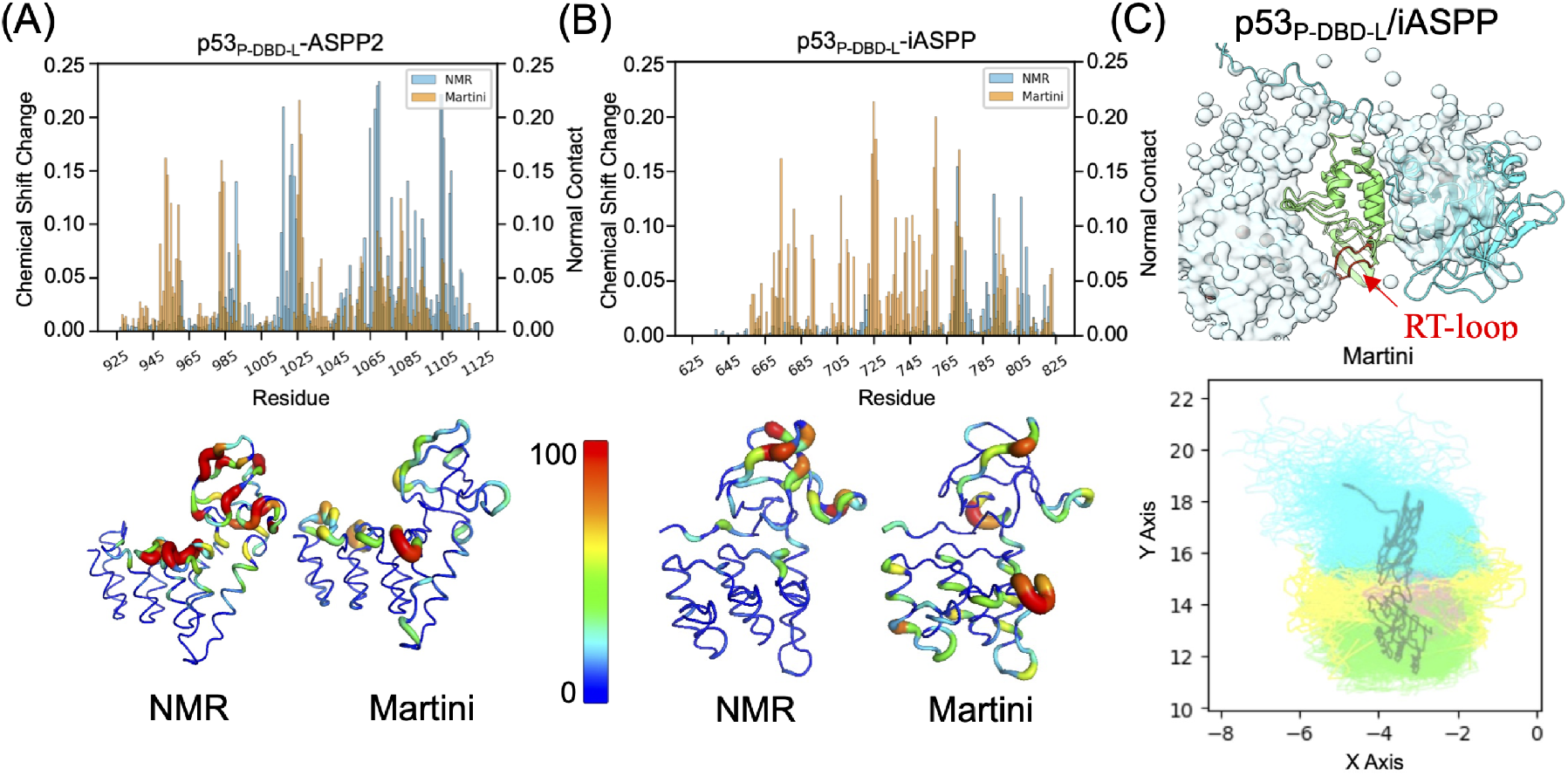
(A-B) Comparison between Martini sampled inter-protein contacts and NMR chemical shifts mapped onto the ANK-SH3 domains of ASPP. NMR is normalized to highest chemical shift change value. Contact scaling is done by summing all contacts involving each ASPP residue and divided by the total number of frames, followed by normalization to C_*max*_ = 0.0002. (C) Martini CGMD sampled of p53_P-DBD-L_-iASPP complexes. The upper panel shows the atoms densities of p53 around ASPP and the lower panel shows the projections of samplings onto the XY plane. The p53’s C-terminal linker (yellow) is interacting with iASPP’s RT (red).

Having validated the Martini CGMD methodology for characterizing PPIs, we proceeded to simulate the binding between p53_P-DBD-L_ and iASPP to investigate whether p53’s C-terminal linker interacts with iASPP’s RT loop, as experimentally demonstrated.^4,43^ We analyzed the atom densities of p53_P-DBD-L_ around iASPP, calculated from the accumulated Martini CGMD trajectories (Fig. 3C), and observed that p53 is positioned around iASPP’s RT loop. Projections of these samplings onto a 2D plane further elucidated the specific interactions. These data provide compelling evidence that the simulated binding complexes accurately capture the interactions between p53’s linker and iASPP’s RT loop, in agreement with experimental findings.

### 3.3 Ensemble-average binding free energy agrees with experimental affinities

We have demonstrated that the ensemble binding complexes qualitatively recapitulate the results of solution NMR and peptide screening assays. Now, we aim to strengthen these outcomes quantitatively. p53DBD binds to the ANK-SH3 domains of ASPP2 and iASPP with comparable affinities (K_*D*_s are 26.4*±* 1.7 nM and 23.3 *±*1.6 nM, respectively^7^). Consequently, we calculated the potential of mean force (PMF) for dissociation and compared it against these experimental K_*D*_s to validate the binding modes.

Firstly, when starting from the crystal-structure-captured single binding mode, iASPP exhibits significantly weaker affinity than ASPP2 in binding p53_DBD_, with binding free energies of*−* 7.57 kcal/mol and *−*12.65 kcal/mol, respectively (Fig. 4A). This discrepancy contradicts the comparable experimental K_*D*_ values. To investigate if starting from multiple binding poses can mitigate this discrepancy, we conducted simulations of the binding process between p53_DBD_ and the ANK-SH3 domains using unbiased Martini CGMD and analyzed the binding modes from various perspectives. Interestingly, for both ASPPs, Martini can capture crystal-like binding modes with RMSD as low as 3.2 Å (Fig. 4B), even though these modes are not the predominant binding modes. To further examine whether the recaptured binding modes are non-specific coincidences resulting from extensive simulation replicas or ASPP-specific binding modes arising from unique protein-protein interactions, we plotted the overall atom density distributions of p53_DBD_ around ASPP (Fig. 4C). This clearly shows that p53_DBD_ exhibits different distribution patterns when bound to ASPP2 and iASPP: Martini-sampled p53_DBD_ conformers are continuously distributed around ASPP2, in contrast to the “bi-sided” distributions around iASPP (Fig. 4C). This suggests that the sampled binding modes are ASPP isoform-dependent. Notably, the “bi-sided” distributions of p53_P-DBD-L_ around iASPP correspond to the major binding modes observed in crystal structures 6RZ3 and 1YCS, implying that iASPP may bind to p53_DBD_ in two predominant modes.

**Figure 4:**
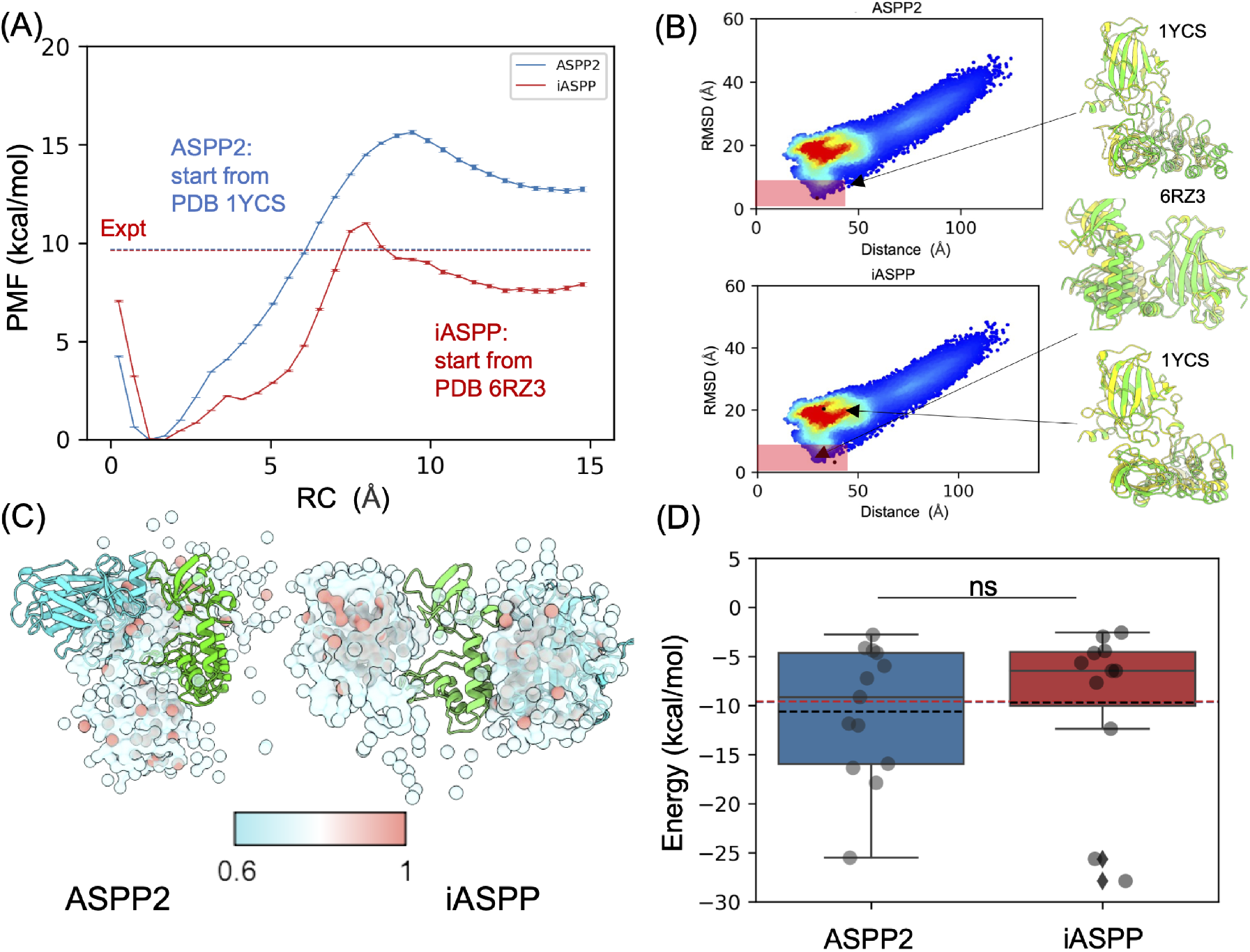
A) PMF curves of the disassociation process between p53_DBD_ and ASPPs, starting from the crystal-structure binding modes. (B) Martini CGMD simulated binding between p53_DBD_ and ASPPs. The trajectories are projected onto the distance (COM of p53_DBD_ to COM of ANKSH3 domain), and the RMSD (with respect to crystal structures) axes. The red shaded areas highlight the binding poses that are close to crystal structures, and those resemble exactly to crystal stuctures are show shown. C) Overall atom density distribution of p53_DBD_ (cyan) around ASPP (green). D) ΔGs calculated from multiple representative binding complexes sampled by Martini CGMD (detailed PMF curves are shown in Fig. S5). The red dashed line corresponds to the experimental ΔG.

Lastly, we selected multiple representative Martini CGMD-sampled p53_DBD_-ASPP complexes and performed umbrella samplings to obtain the dissociation PMFs. We employed a rigorous selection procedure to maximize the representativeness of the selected binding complexes (see Section Sect. S2). As shown in Fig. 4D, ASPP2 exhibits different distributions of binding free energies compared to iASPP: ASPP2 includes several cases with weak binding, which are compensated by tight binding ones, while iASPP cases are more concentrated around the experimental K_*D*_s. However, the average values of calculated binding free energies show no statistically significant difference (P = 0.995, t-test). Calculated binding free energies starting from crystal structures do not align with the experimental observations that iASPP and ASPP2 have comparable binding affinities toward p53_DBD_. Instead, the ensemble-average energies based on Martini CGMD-sampled multiple binding poses align with the experimental observations.

### 3.4 p53’s IDRs accelerate the binding rates and further differentiates p53’s bindings toward ASPP2 and iASPP

We further investigated the impact of the intrinsically disordered regions (IDRs) in the p53 protein, which constitute approximately 48% of the entire protein (Fig. S1), on the PPIs based on our Martini CG trajectories. Comparison between p53_DBD_ and p53_P-DBD-L_ shows that addition of IDRs to p53_DBD_ significantly reduces the mean first passage time for inter-protein contact formation, from approximately 130 ns to approximately 90 ns (Fig. 5A). This indicates an enhancement in the association rates facilitated by the presence of IDRs. Interestingly, while the addition of IDRs to p53 does not significantly alter the kinetics of binding to ASPP2 and iASPP, it does influence the specific inter-protein contacts in the simulated complexes. We calculated the time-dependent and p53-domain-specific inter-protein contacts (Fig. 5B). These results clearly demonstrate that the presence of p53’s IDRs leads to distinct interactions: When binding to ASPP2, p53’s DBD domain maintains considerable contacts, whereas for binding to iASPP, the interaction relies less on the DBD domain and is primarily mediated by p53’s linker (Fig. 5B).

**Figure 5:**
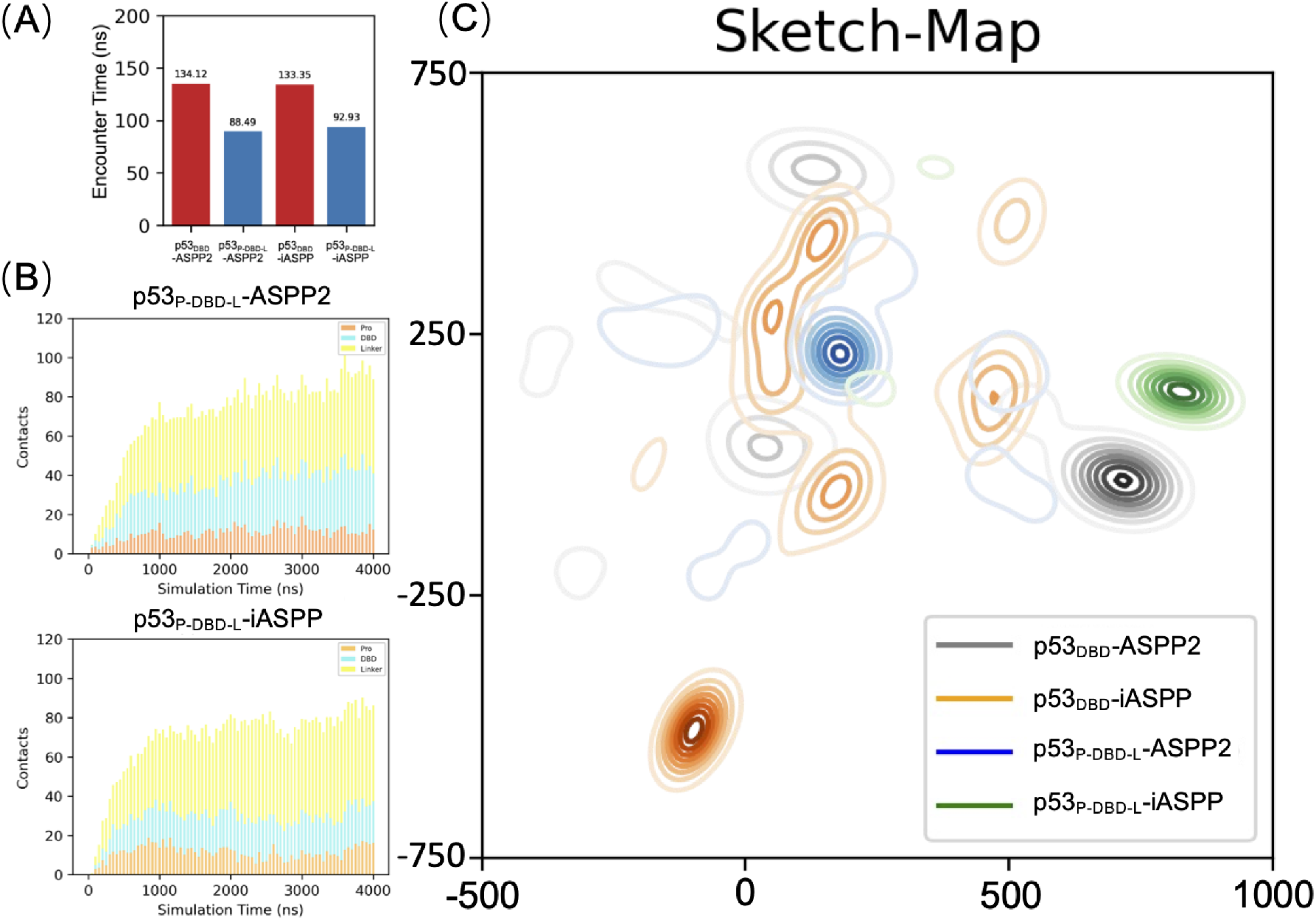
A) Average encounter time when the first contact between p53 and ASPP occurs. B) Average time-dependent number of contacts between ASPP and different p53 domains calculated from 50*×* 4 µs Martini CG simulations. C) Projections of Martini CGMD sampled ASPP-p53 complexes onto 2D plane using the sketch-map dimentionality reduction method.

Furthermore, the presence of p53’s linker serves to differentiate the binding patterns between iASPP and ASPP2 when interacting with p53. To illustrate this, we employed the sketch-map dimensionality reduction method to project the simulated ASPP-p53 complexes onto a 2D plane. The choice of sketch-map enables precise projection of the relative configurations of ASPP-p53 complexes in 3D space onto the 2D plane. Consequently, points that are close in the 2D plane represent similar ASPP-p53 binding configurations, allowing for straightforward comparison of the simulated ASPP-p53 complexes. As shown in Fig. 5C, in the absence of IDRs, the binding of p53_DBD_ to iASPP2 and ASPP2 exhibits moderate overlap, despite differing major binding poses (i.e., high-density areas differ for iASPP2 and ASPP2). Notably, the addition of IDRs to p53_DBD_ leads to a dramatic change in the distribution patterns for both iASPP and ASPP2 on the 2D projection plane. This results in more concentrated and distinct binding patterns for these two ASPPs. These projection data strongly suggest that the inclusion of p53’s IDRs further differentiates p53’s binding interactions with ASPP2 and iASPP.

## 4 Discussion

Two lines of evidences suggested that the iASPP-p53 binding mode may be different from that resolved in PDB 6RZ3, and this alternative mode is highly similar to the ASPP2-p53 binding mode resolved in PDB 1YCS. Firstly, the isolated iASPP ANK-SH3 crystal structure (PDB 2VGE^7^) was found to use a surface patch, which resembles the ASPP2’s binding patch towards p53, to interact with neighbouring unit. Secondly, solution NMR data shows that the p53-induced residue chemical shift perturbations were mapped onto the similar surfaces for ASPP2 and iASPP ANK-SH3 domains.^4^ The ongoing debate over the binding mode presents challenges to achieving a comprehensive understanding of ASPP-p53 regulations.

Discrepancies between NMR and crystallography-determined modes are not uncommon in the literature, not only for individual proteins but also for protein-protein interactions (PPIs). While NMR and crystallography-derived protein structures align closely in most cases,^12^ certain instances can exhibit substantial structural deviations, with backbone RMSD reaching up to 6 Å.^13^ This divergence can be attributed to the distinct environments in which the structures are determined, namely, aqueous solution versus crystal lattice. Regarding PPIs, assigning solution NMR-determined PPIs as biologically relevant protein binding modes is generally less contentious. However, the same cannot be said for crystallography-determined PPIs due to the influence of crystal packing. Crystal packing effects can introduce free energy advantages that compete with native PPI formations.^11,45,46^Consequently, the relevance of crystallography-determined PPIs to biological functional complexes is affinity-dependent.^45^ Strong PPIs (e.g., K_*D*_ <100 µ M^45^) are more likely to retain their native binding modes within crystal lattices. In our specific case, the binding affinities between p53’s DBD and ASPP’s ANK-SH3 domains are around the µM range as measured by different methods,^7,4,8^which is in proximity to the 100 µM threshold. This brings uncertainty in assuring the crystal structure captured ASPP-p53 complexes as biological-relevant assembly, especially for the iASPP-p53 PPI.

Our simulated p53-ASPP ensemble binding complexes unify the and crystallography- and NMR- modes. Firstly, we demonstrated that the ensemble of binding complexes better aligns with experimental observations, supported by two key arguments: 1) the ensemble binding modes satisfy the experimental-confirmed binding patterns while the single crystal structure does not. 2) Calculated binding free energies, using representative complexes from the ensemble, closely match the experimental K_*D*_ values. More importantly, the simulated complexes contain the exact crystal-structure binding modes, and for iASPP, the simulated complexes exhibit “bi-sided” distributions of p53 around the ANK-SH3 domain, which can be treated as two generalized binding modes representing the NMR-suggested and crystallography-captured binding mode, respectively. Therefore, p53_DBD_ binds to iASSP’s ANK-SH3 domain in two configurations in solution, and crystal packing shifts the binding equilibrium to the one captured in PDB 6RZ3. This potentially elucidates why iASPP utilizes similar binding modes as ASPP2 towards p53, as suggested by solution NMR, while exhibiting a different binding mode in the crystal structure. As a result, our simulated ASPP-p53 complexes are complementary to the NMR and crystallography characterizations of the p53-ASPP complexes.

IDRs are not merely flexible but have envolved to exert unique functions such as allowing rapid and reversal PPI and acting as recognition motifs.^47,48^ Our simulation data suggest that p53’s IDRs play multiple roles in fine-tuning the PPI. IDPs are well-known to accelerate protein binding rates through the “fly-casting” mechanism^49^ which states that dynamic IDP increases the searching radius to capture binding partner. This was reflected in our simulations as we show that adding the N-terminal Pro-domain and C-terminal linker to p53_DBD_ greatly accelerates the binding to both ASPP and iASSP. While the rate-accelerating effect brought by p53’s IDRs are similar between ASPP2 and iASPP, the simulated protein-protein binding complexes (e.g., the encounter complexes^50^) are differently impacted by p53’s IDRs. To visualize the difference, we elected to use the sketch-map dimensionality reduction method to project the ASPP-p53 complex conformation onto the 2D plane. We show that adding IDRs to p53_DBD_ significantly reduced the overlap between ASPP2 and iASPP, indicating more different samplings..

One angle to explain the functional consequence of differently-sampled ASPP-p53 encounter complexes is through the concept of “effective concentration”.^51,52^ Protein-protein binding process can be divided into two sub-processes:^44^ 1) Diffusion of protein partners to form loosely bound complexes (encounter complexes). 2) Rearrangement of encounter complex to reach native PPI state, such as rolling on another protein’s surface.^53^ While step one is driven by diffusion, step two (refolding) is conditional, and occurs on subsets of the encounter complex that satisfy certain criteria such as shape complementary.^54^ In this regard, adding IDRs to p53 increase the size of the candidate pool (effective concentration) that from which more complexes can refold to the native PPI state.

However, the mechanism by which ensemble binding of PPIs exerts its biological functions remains elusive, and addressing this question requires further structural characterizations encompassing the entire ASPP and p53 proteins. Our argument is supported by several studies demonstrating that motifs distant from the folded domains contribute to their biological function. For example, peptide screening assays have shown that ASPP2’s Pro-domain can autoinhibit the ANK-SH3 domain, competing with p53’s binding.^47^ Additionally, an ASPP2 isoform with the N-terminal region deleted inhibits p53-dependent apoptosis, exhibiting a completely different regulatory effect on p53 compared to full-length ASPP2.^55^ Even for ASPP1 and ASPP2 proteins, both of which promote p53-dependent apoptosis and have nearly identical sequences and folds in their ANK-SH3 domains, there is a *∼*10-fold difference in their affinities for p53, with K_*D*_=5 µM for ASPP2 and K_*D*_=0.5 µM for ASPP1, respectively. However, ASPP2 can displace the DNA promoter from the p53-DNA promoter complex, while ASPP1 cannot.^9^ This contradiction, where ASPP2 has weaker affinity with p53 but can still displace the DNA promoter, implies that other parts of ASPP are involved in *in vivo* regulation. Therefore, a comprehensive understanding of the contrasting regulatory effects of ASPP2 and iASPP on p53 requires structural and biological investigations at the whole protein level in PPIs.

## 5 Conclusions

In this study, we performed multi-scale MDs and free energy calculations to characterize the PPI of two proteins that of great therapeutic potentials, the p53 and its regulator(s), iASPP/ASPP2. Our work focus on reconciling the discrepancy between NMR- and crystallography- determined binding modes for the iASPP-p53 PPI. We used unbiased Martini3.0 CGMD to simulated the proteinprotein PPI, and achieved the ASPP2 and iASPP-specific ensemble binding modes that can unify the reported binding mode discrepancy. We first validated the ensemble binding modes by showing that the ensemble-average inter-protein contacts are well aligned with solution NMR-detected chemical shift perturbations caused by p53. Furthermore, the ensemble-average PPI binding free energies agree with experimental Kds. Notably, we observed that the crystallography-determined binding modes were recapitulated within the ensemble-binding modes, indicating that the crystal packing environment may influence the equilibrium of binding modes toward the configuration captured by crystallography. Although linking these ensemble binding modes to actual biological functions poses a challenge, we provided evidence that the ensemble binding modes are highly sensitive to p53’s intrinsically disordered regions (IDRs). The inclusion of IDRs accentuated the differences in binding modes exhibited by p53 towards ASPP2 and iASPP. This sensitivity underscores the biological significance of ensemble bindings and is consistent with observations indicating that the entire proteins, rather than just their folded domains, contribute to biological functions.

## 6 Acknowledgements

Research reported in this work was supported by the Harbin Medical University high-level introduction of talent research start-up fund (No.310212000109, 31011210004) and the National Science Foundation (No. 22002096).

## Supplementary Information (SI)

**Table S1:**
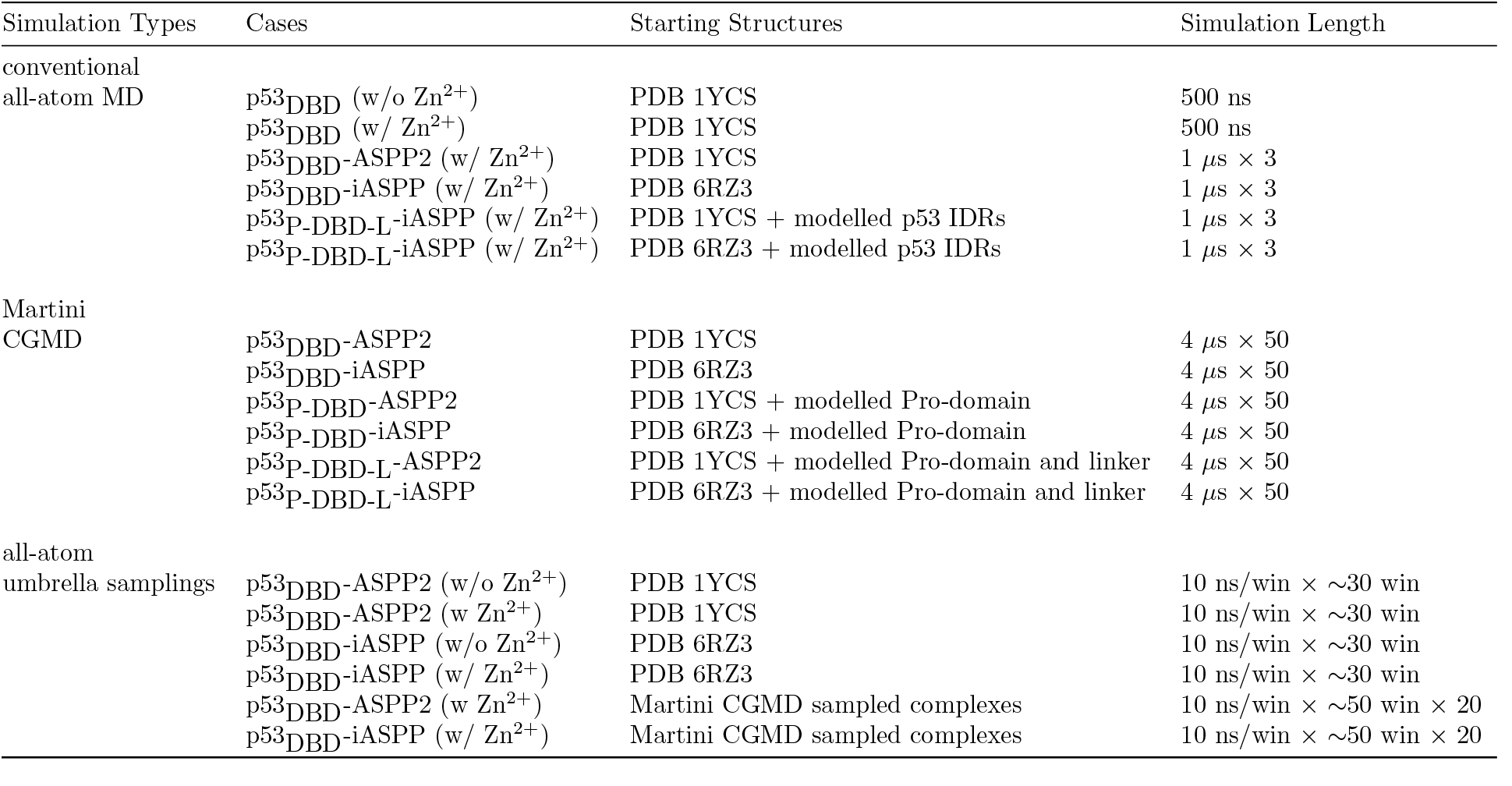
Molecular dynamics simulations performed in this study.

**Figure S1:**
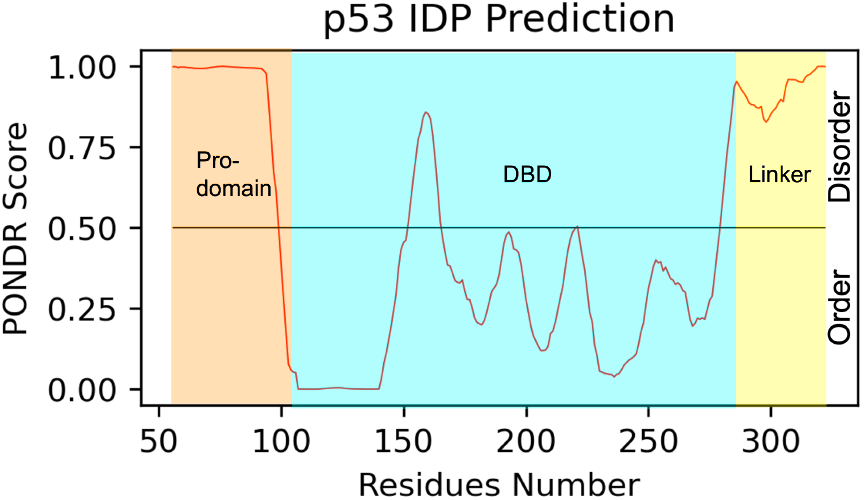
The intrinsically disordered domains of p53 as predicted by the online-website PONDR [56]. The Pro-domain and linker-domain flanking p53_DBD_ are disordered.

## S1 Zn^2+^ plays a critical role in stabilizing p53_DBD_ and in affecting the binding free energy between p53_DBD_ and ANK-SH3 domain of ASPP proteins

A Zn^2+^ cation was resolved in p53_DBD_ crystal structure (PDB 1YCS) to coordinate with p53’s His83-CYS80,142,146 motif. To investigate the importance of Zn^2+^, we performed MD simulations on p53_DBD_ with and without the Zn^2+^, respectively. We found that removing Zn^2+^ leads to significant conformational change of p53_DBD_ and alters the dynamics of the domain, as compared to the w/ Zn^2+^ case (Fig. S2A-B). Specifically, the L3 loop of p53_DBD_ is distorted after removing the Zn^2+^. This loop is at the PPI interface to bind ASPP2 (PDB 1YCS) [47], and has been shown to hub most cancer-causing mutations [57]. Therefore, Zn^2+^ role of stabilizing this loop needs to be considered. We also investigate the role of Zn^2+^ in affecting the potential of mean force (PMF) along the p53_DBD_-ANK-SH3 disasscotion pathway. As shown in Fig. S2C-D, presence of Zn^2+^ can significantly change the PMF curves for both ASPP2 and iASPP.

**Figure S2:**
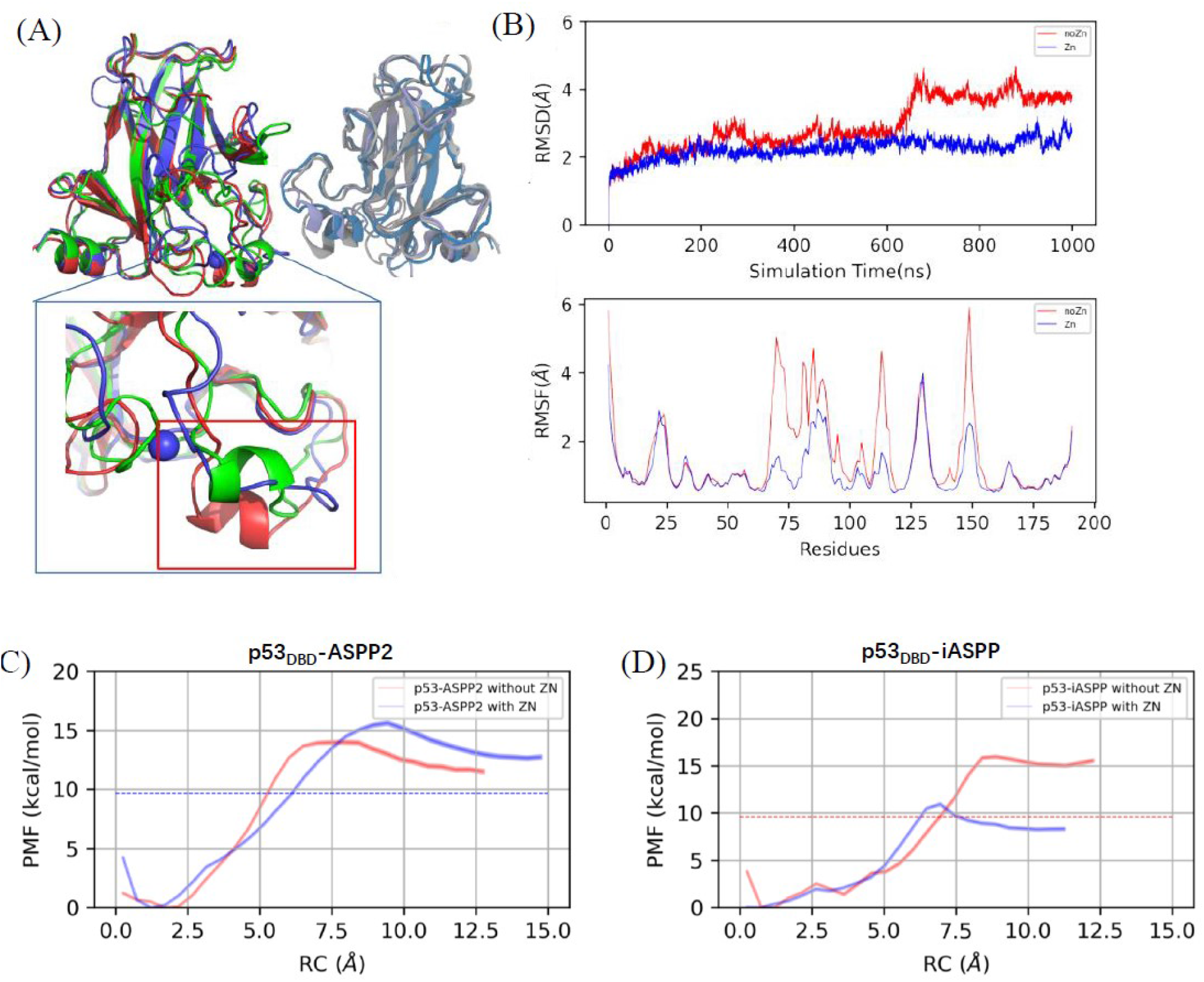
(A) Superimposing MD-sampled p53_DBD_ conformations, w/ Zn^2+^ (blue) and w/o Zn^2+^ (red), on crystal structure (green). (B) RMSD and RMSF of p53_DBD_ from MD simulations w/ and w/o Zn^2+^. (C-D) PMF of p53_DBD_ disassociation from ASPP2 and iASPP w/ and w/o Zn^2+^.

## S2 Selecting representative p53-ASPP complexes from Martini CGMD sampled protein-protein complexes

A center of mass (COM) distance criteria (<30 Å) and RMSD criteria (<50 Å, with respect to PDB 1YCS) were applied on the accumulated Martini trajectories to filter out frames that do not have p53 and ASPP contacted. The surviving trajectory frames were then aligned on p53_DBD_ and were subject to following processes: ASPP protein densities around p53_DBD_ were calculated using the *grid* command from the CPPTRAJ program. Regions have high relative density (> 0.6) were identified as the most probable binding patterns. The COM of ASPP was drawn around p53 for each frame. For those frames their COMs are located within the high density regions, they are identified as candidates. 20 frames were randomly picked out from the candidate pool.

**Figure S3:**
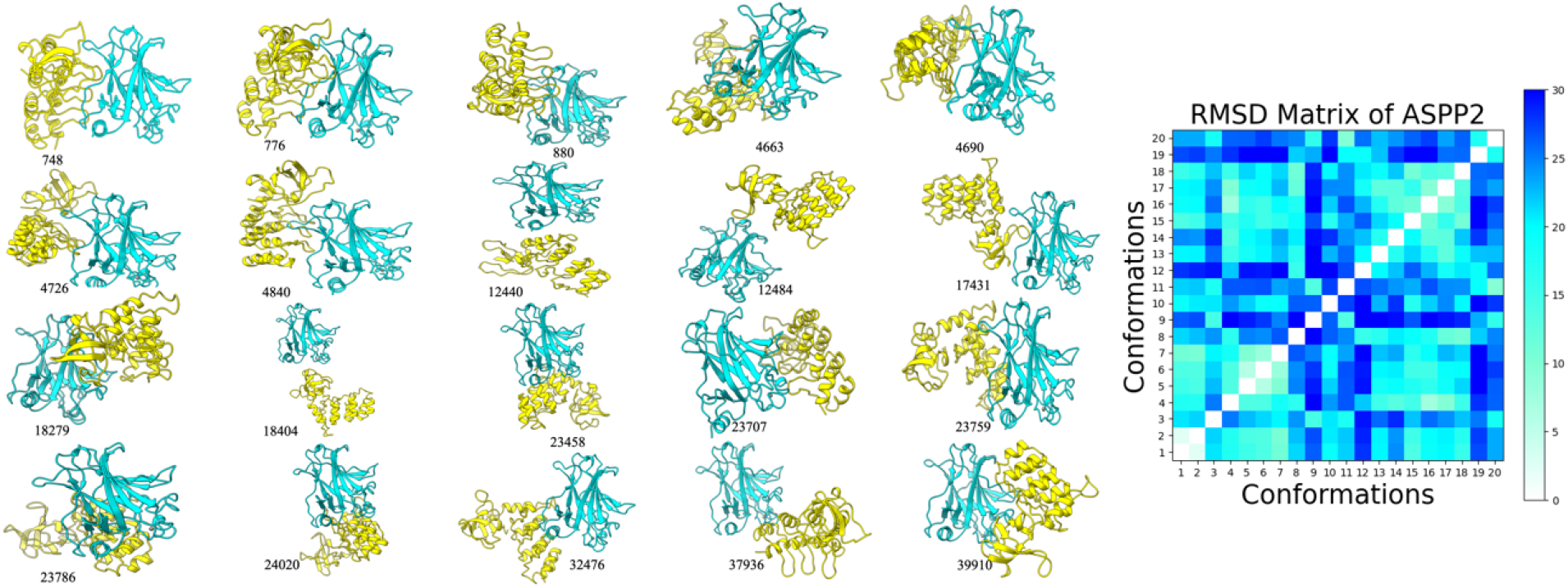
20 representative p53_DBD_-ASPP2 complexes sampled by Martini CGMD. The pairwise RMSDs of ASPP2 (after aligning on p53) were shown on the right to illustrate the structural difference of ASPP2.

**Figure S4:**
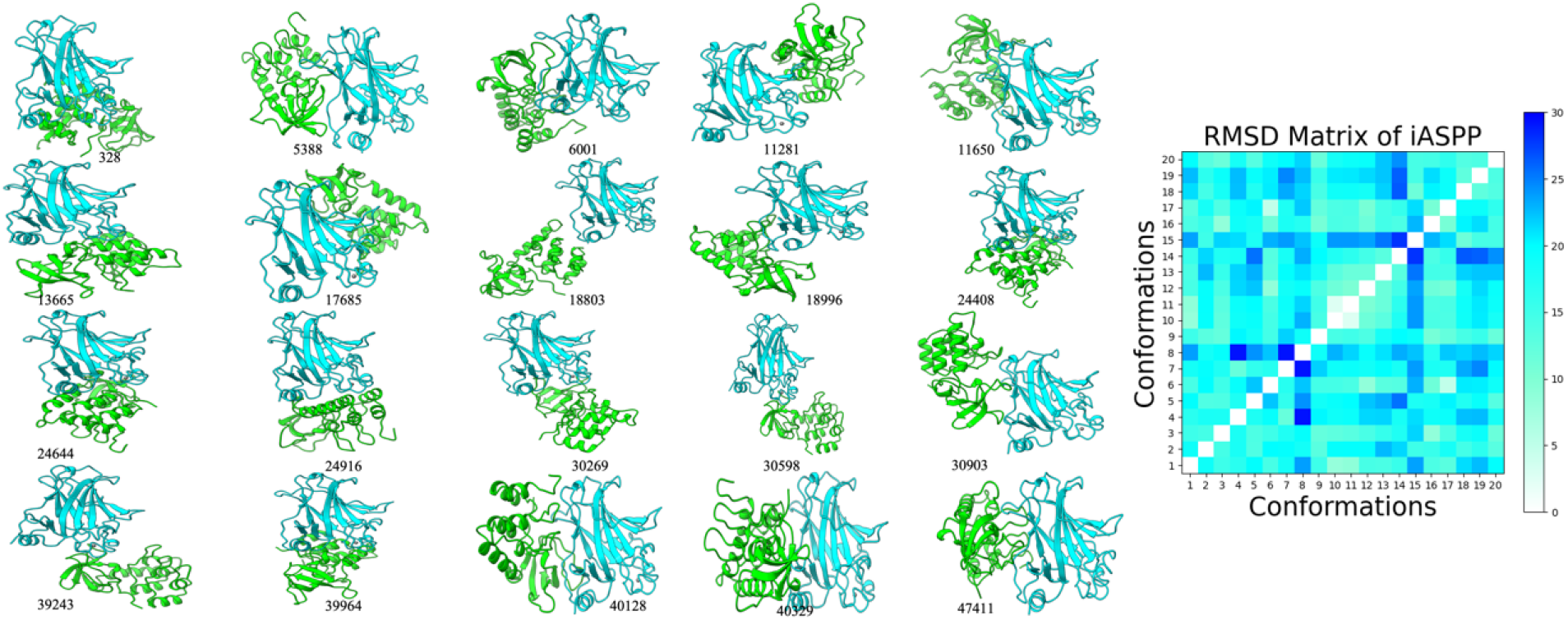
20 representative p53_DBD_-iASPP complexes sampled by Martini CGMD. The pairwise RMSDs of iASPP (after aligning on p53) were shown on the right to illustrate the structural difference of iASPP.

**Figure S5:**
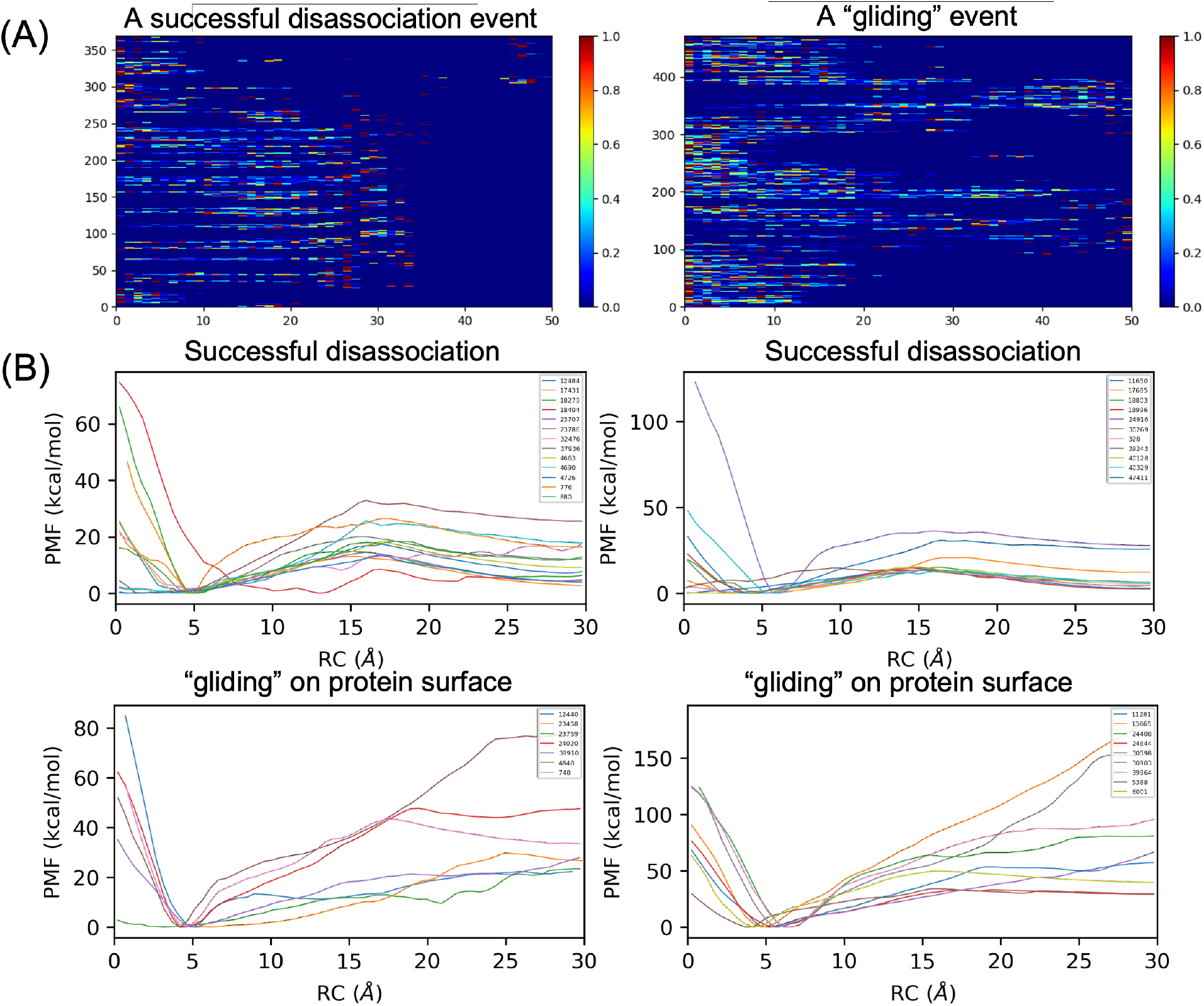
(A) Umbrella sampling using COM-COM distance as CV can lead to protein gliding on another protein’s surface instead of driving complex directly disassociate. We show that during the umbrella sampling, a successful disassociation has inter-protein contacts smoothly disappear as window number increases, indicating a clean one-way disassociation, while for a “gliding” event, newly formed inter-protein contacts are keep emerging as window number increases. (B-C) For the 20 representative Martini CGMD-sampled complexes for each ASPP protein, the PMF curves for successful disassociations (n=13/20 for ASPP2, and n=11/20 for iASPP), and for “glidings” are given.

## Notes

### Competing Interest Statement

The authors have declared no competing interest.

